# Network efficiency predicts resilience to cognitive decline in elderly at risk for Alzheimer’s

**DOI:** 10.1101/2020.02.14.949826

**Authors:** Florian U. Fischer, Dominik Wolf, Andreas Fellgiebel, for the Alzheimer’s Disease Neuroimaging Initiative

## Abstract

To determine whether white matter network efficiency (WMNE) may be a surrogate marker of the physiological basis of resilience to cognitive decline in elderly persons without dementia and age and AD-related cerebral pathology, we quantified WMNE from baseline MRI scans and investigated its association with longitudinal neuropsychological assessments independent of baseline amyloid, tau and white matter hyperintensity volume. 85 cognitively normal elderly subjects and patients with mild cognitive impairment (MCI) with baseline diffusion imaging, CSF specimens, AV45-PET and longitudinal cognitive assessments were included. WMNE was calculated from reconstructed cerebral white matter networks for each individual. Mixed linear effects models were estimated to investigate the association of higher resilience to cognitive decline with higher WMNE and the modulation of this association by increased cerebral amyloid, CSF tau or WMHV. For the majority of cognitive outcome measures, higher WMNE was associated with higher resilience to cognitive decline independently of pathology measures (beta: .074 – .098; p: .011 – .039). Additionally, WMNE was consistently associated with higher resilience to cognitive decline in subjects with higher cerebral amyloid burden (beta: .024 – .276; p: .000 – .036) and with lower CSF tau (beta: −.030 – −.074; p: .015 – .002) across all cognitive outcome measures. The results of this study indicate that WMNE in particular and possibly white matter organization in general may be worthy targets of investigation to provide measures quantifying a patient’s resilience to cognitive decline and thus provide an individual prognosis.

## Introduction

In-vivo amyloid imaging has profoundly improved the diagnosis of Alzheimer’s disease (AD) at its pre-dementia stages. However, individual predictions of cognitive decline are unsatisfactory from a clinical perspective due to considerable variance (Donohue et al., 2017; Insel et al., 2016; Jack et al., 2016; Vos et al., 2013), which presumably refers to an individual’s capacity to tolerate or compensate cerebral pathology, commonly termed reserve or – more generally – resilience (Barulli & Stern, 2013; Cabeza et al., 2018; Wolf, Fischer, & Fellgiebel, 2018; Yaffe et al., 2011). The identification and quantification of an MRI-based surrogate of the physiological basis of this resilience could thus complement and significantly improve individual predictions of cognitive decline based on cerebral pathology. However, the underlying physiological basis of resilience to cognitive decline has not conclusively been identified so far. In this regard, white matter (WM) has received surprisingly little attention, although WM microstructural properties have repeatedly been demonstrated to be associated across the lifespan, including old age, with intelligence (Bathelt, Scerif, Nobre, & Astle, 2018; Fischer, Wolf, Scheurich, & Fellgiebel, 2014; Koenis et al., 2018; Li et al., 2009; Schmand, Smit, Geerlings, & Lindeboom, 1997), which is a known factor of resilience to neurodegenerative disease (Stern, 2012; Whalley, Deary, Appleton, & Starr, 2004). Furthermore, functional imaging studies have demonstrated that maintained functional connectivity is more associated with cognitive performance at higher age (Tsvetanov et al., 2016, 2018) and that the amount of neural resources that are recruited to process cognitive tasks is increased in higher age and in the presence of cerebral pathology (Reuter-Lorenz & Park, 2014; Stargardt, Swaab, & Bossers, 2015). As WM facilitates information transfer and synchronization between networks of spatially distributed grey matter (GM) regions that are functionally joined in the processing of more demanding cognitive tasks, one might expect that higher WMNE modulates the efficacy of the integration and indeed, WMNE is associated with intelligence across the lifespan (Bathelt et al., 2018; Fischer et al., 2014; Li et al., 2009). For these reasons we hypothesized that WMNE may be a candidate surrogate measure of resilience to cognitive decline.

## Methods

Data used in the preparation of this article were obtained from the Alzheimer’s Disease Neuroimaging Initiative (ADNI) database (adni.loni.usc.edu). The ADNI was launched in 2003 as a public-private partnership, led by Principal Investigator Michael W. Weiner, MD. The primary goal of ADNI has been to test whether serial magnetic resonance imaging (MRI), positron emission tomography (PET), other biological markers, and clinical and neuropsychological assessment can be combined to measure the progression of mild cognitive impairment (MCI) and early Alzheimer’s disease (AD). For up-to-date information, see www.adni-info.org.

### Subjects

Subjects and their respective data points were selected from the database of the ADNI project according to the following criteria: baseline assessment during the ADNI 2 phase and classification as cognitively normal or MCI, availability of T1, FLAIR, diffusion weighted (DWI) MRI as well as florbetapir (AV45) PET imaging at baseline as well as longitudinal neuropsychological assessments. MCI subjects were included so as to ensure sufficient variance of cognition and pathological markers in a continuum of elderly non-demented subjects for the assessment of possible resilience. For details regarding the cognitive assessment within the ADNI, please refer to the publicly available procedures manual: https://adni.loni.usc.edu/wp-content/uploads/2008/07/adni2-procedures-manual.pdf. In total, 85 subjects consisting of 34 females and 51 males aged 56.5 to 89.0 years were included (see table 1 for details descriptive sample characteristics).

**Table 1:**
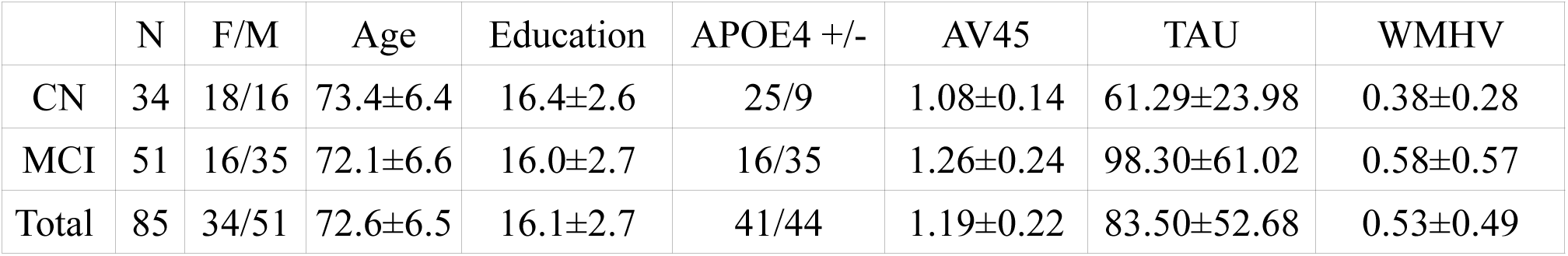
Descriptive statistics of sample demographics and cerebral pathology measures. CN: cognitively normal. MCI: mild cognitive impairment. APOE4: apolipoprotein ε4. AV45: Global4. AV45: Global florbetapir standardized uptake value ratio. TAU: liquor specimen total tau. WMHV: normalized WMHV in percent.

|  | N | F/M | Age | Education | APOE4 +/- | AV45 | TAU | WMHV |
| --- | --- | --- | --- | --- | --- | --- | --- | --- |
| CN | 34 | 18/16 | 73.4±6.4 | 16.4±2.6 | 25/9 | 1.08±0.14 | 61.29±23.98 | 0.38±0.28 |
| MCI | 51 | 16/35 | 72.1±6.6 | 16.0±2.7 | 16/35 | 1.26±0.24 | 98.30±61.02 | 0.58±0.57 |
| Total | 85 | 34/51 | 72.6±6.5 | 16.1±2.7 | 41/44 | 1.19±0.22 | 83.50±52.68 | 0.53±0.49 |

### CSF measurement

All CSF biomarkers collected at different centers were stored and analyzed at the Penn ADNI Biomarker Core Laboratory at the University of Pennsylvania, Philadelphia. CSF concentrations of total tau were measured in the baseline CSF samples using the multiplex xMAP Luminex platform (Lumnix Corp., Austin, TX). We included total tau instead of phosphorylated tau, as the latter is included in the former and also more specific to AD and thus limited in scope to AD-specific tauopathy (Blennow, Vanmechelen, & Hampel, 2001). More details on data collection and processing of the CSF samples can be found elsewhere (Shaw et al., 2009) (http://adni.loni.usc.edu/methods).

### APOE genotype

Apolipoprotein (APOE) genotype was determined by genotyping the two single nucleotide polymorphisms that define the APOE ε2, ε3, and ε4 alleles (rs429358, rs7412) with DNA extracted2, ε2, ε3, and ε4 alleles (rs429358, rs7412) with DNA extracted3, and ε2, ε3, and ε4 alleles (rs429358, rs7412) with DNA extracted4 alleles (rs429358, rs7412) with DNA extracted by Cogenics from a 3-ml aliquot of EDTA blood (http://adni.loni.usc.edu/data-samples/genetic-data).

### Imaging data acquisition

DWI, FLAIR and inversion-recovery spoiled gradient recalled (IR-SPGR) T1-weighted imaging data were acquired on several General Electric 3T scanners using scanner specific protocols. Briefly, DWI data was acquired with a voxel size of 1.37^2^ × 2.70 mm^3^, 41 diffusion gradients and a b-value of 1000 s/mm^2^. IR-SPGR data were acquired with a voxel size of 1.02^2^×1.20mm^3^. AV45 PET imaging data were acquired on several types of scanners using different acquisition protocols. To increase data uniformity, the data underwent a standardized preprocessing procedure at the ADNI project. All imaging protocols and preprocessing procedures are available at the ADNI website (http://adni.loni.usc.edu/methods/).

### T1-weighted and FLAIR data processing

The T1-weighted IR-SPGR data was automatically tissue segmented and spatially normalized to MNI-space using SPM8 (http://www.fil.ion.ucl.ac.uk/spm/) and the VBM8-toolbox (http://www.neuro.uni-jena.de/vbm/). Additionally, inverse transformations from MNI to native T1 space were calculated.

Grey matter (GM) was segmented into 106 functionally and anatomically defined cortical regions as well as the subcortical basal ganglia regions as implemented in the probabilistic Harvard Oxford Atlas supplied with FSL.

TIV as well as hyperintensity volume (WMHV) were calculated at ADNI core laboratories from T1-weighted and FLAIR data using published tissue segmentation methods (DeCarli, Fletcher, Ramey, Harvey, & Jagust, 2005; Fletcher, Singh, Harvey, Carmichael, & DeCarli, 2012). WMHV was also normalized dividing by the TIV.

### DWI data processing

DWI data were corrected for eddy currents and motion artefacts using the method of Rohde et al. as implemented in VistaSoft (Rohde, Barnett, Basser, Marenco, & Pierpaoli, 2004); diffusion gradients were adjusted according to the resulting transformations. Additionally, DWI data were upsampled to 1mm isotropic voxel size for further processing. For fiber tractography, Anatomically Constrained Tractography (ACT) as implemented in MRtrix was employed (Smith, Tournier, Calamante, & Connelly, 2012). This approach incorporates anatomical constraints based on tissue segmentations of T1 data. To this end, VBM8 tissue segmented data in T1 native space were coregistered using SPM8 to the upsampled DWI B0 images, and used in the subsequent ACT. Based on the tissue segmentation images, the ACT framework calculates an isocontour representing the interface of GM and WM for fiber seeding. Subsequently, tractography seed points were placed randomly along the GM-WM interface. Starting from these points, the probabilistic “ifod2” tractography algorithm was executed until 500,000 anatomically plausible streamlines were reconstructed for each subject. Streamlines were accepted if they met the anatomical constraints of ACT (Smith et al., 2012).

### Network reconstruction and characterisation

In order to reconstruct fibers, the Harvard Oxford Atlas GM ROIs were first warped to native T1 space using the inverse VBM8 normalization transformations and then transferred to the upsampled DWI space using the transformation estimated from the T1 to B0 coregistration.

Subsequently, for each ROI pair in each subject, the number of connecting streamlines was obtained and recorded to construct the adjacency matrix. A connection threshold of at least 3 connecting streamlines for each connection was applied (Fischer et al., 2014; Li et al., 2009). Finally, WMNE for each subject was calculated as the average efficiency of the network according to the formula

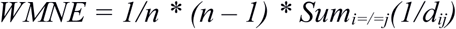

where *n* represents the number of nodes and *d*_*ij*_ is the inverse of the connection weights (Latora & Marchiori, 2001). For a detailed description and discussion of graph measures see (Fornito, Zalesky, & Breakspear, 2015).

### PET data processing

Subjects’ global cortical amyloid-β load was calculated from florbetapir (AV45) PET images according to procedures established by the ADNI (http://adni.loni.usc.edu/methods/pet-analysis/). Briefly, cortical amyloid was calculated as the average of the AV45 uptake in the frontal, angular/posterior cingulate, lateral parietal and temporal cortices normalized by dividing by the mean uptake in the cerebellum.

### Neuropsychological assessment and quantification of resilience

Subjects underwent an extensive neuropsychological assessment. For details regarding the cognitive assessment within the ADNI, please refer to the publicly available procedures manual: https://adni.loni.usc.edu/wp-content/uploads/2008/07/adni2-procedures-manual.pdf. Within the scope of this study, the cognitive Alzheimer’s Disease Assessment Scale (ADAS-cog), which spans several cognitive domains (Rosen, Mohs, & Davis, 1984), as well as well as the Clinical Dementia Rating Scale Sum of Boxes (CDRSOB) and the Minimental State Examination (MMSE) were investigated.

### Approach to studying resilience mechanisms

The resilience modeling approach used in this paper has been published and discussed previously (Fischer, Wolf, & Fellgiebel, 2019; Wolf et al., 2018). For an extensive description, please refer also to the supplement. In brief, resilience can very generally be defined as cognitive outcome variance unexplained by pathology, which can be statistically modelled by partialling associations with pathology out of the cognitive outcome variable. The remaining variance, termed resilience, can be associated with the resilience factor, which is termed *general resilience*. However, pathology might modulate this association, which is termed *dynamic resilience* in the case of a positive interaction and *limited resilience* in the case of a negative interaction. To accommodate longitudinal data, a time variable can be included, as well as its interaction terms with pathology measures, the resilience factor and their interaction term:

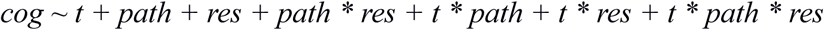

where *cog* represents cognitive outcome, *t* the time variable, *path* pathology measures and *res* the resilience factor.

### Statistical analysis

Longitudinal ADAS-cog, CDRSOB and MMSE measures were coded in single variables and set as cognitive outcome and thus as the dependent variables. The time of assessment of the cognitive outcome variables was encoded as an additional time variable that was set as an independent variable. Other independent variables consisted of baseline WMNE as potential resilience factor as well as AV45, TAU and WMHV as measures of baseline pathology. For each of these, a family of models were estimated representing all possible combinations of the following terms: WMNE, the interaction term of WMNE and the time variable, the first order interaction terms of WMNE and pathology measures as well as the second order interaction terms of WMNE, the time variable and pathology measures. Note that for models including interaction terms, the constituent terms were also always included in the models - including all three possible first order interaction terms of the individual elements of second order interaction terms. In total, this yielded 36 models to be estimated for each cognitive outcome measure. For a list containing the terms of each model, please refer to the supplement.

The models were estimated as linear mixed effects models, wherein random intercepts for each subject and for clinical status at baseline (CN/MCI) were estimated. Additionally, all models included the baseline pathology measures AV45, TAU and WMHV as well as their interaction terms with the time variable. Years of education, age, gender and APOE4 positivity were included in all models as covariates. Subsequently, for each cognitive outcome measure and the corresponding family of 36 possible resilience models, the conditional Akaike Information Criterion (cAIC)was calculated (Greven & Kneib, 2010) as well as the Delta cAIC scores (differences between the model with the lowest cAIC score and each of the remaining models), the Akaike weights and the marginal coefficient of determination. All models with Delta cAIC lower than 2 were considered as models with substantial evidence.

The model terms of interest as per the extended resilience model described in the previous section were the interaction term between the time variable and WMNE (general resilience to cognitive decline) as well as the second order interaction terms of the time variable with WMNE and with baseline pathology measures (dynamic or limited resilience to cognitive decline). For these terms, significance testing was carried out if they were included in all models with Delta cAIC lower than 2 for each cognitive outcome. Significance testing was conducted using the LRT test as well as parametric bootstrapping at 1 million simulations per test. In the case of dynamic or limited resilience, the model with the lowest cAIC was compared to a reduced model with the term of interest (*t* * *WMNE* * *path*) removed. In the case of general resilience to cognitive decline, different models had to be compared, as the term of interest (*t* * *WMNE*) is a constituent term of all second order interaction terms. For this reason, all second order interaction terms were removed from the model with the lowest cAIC. This model was then compared to a further reduced model with the term of interest removed. Furthermore, the models with lowest cAIC were reestimated using robust linear mixed regression models.

All statistical analyses were conducted using R 3.4.0 as well as the packages “lme4” (Bates, Mächler, Bolker, & Walker, 2014) “robustlmm” (Koller, 2016), “cAIC4” (Säfken, Rügamer, Kneib, & Greven, 2018), “pbkrtest” (Halekoh & Højsgaard, 2014) as well as “car” (Fox & Weisberg, 2019). For cAIC and significance testing, models were estimated with maximum likelihood estimation. For coefficient reporting, models were reestimated using restricted maximum likelihood estimation. Two data points with implausibly low MMSE measurements (4 and 6) were removed in the respective models. All variables that were part of an interaction term in any model were centered to reduce collinearity (Aiken, West, & Reno, 1991). Variance inflation was calculated for the best model for each cognitive outcome variable. WMHV was log-transformed to achieve approximate normal distribution. The significance threshold was set to α = 0.05 for all analyses. Results were corrected for FDR at 5%.

## Results

The models with lowest cAIC score were (31) for ADAS-cog, (34) for CDRSOB and (31) for MMSE (see supplement). For ADAS-cog and MMSE, model (10) showed a Delta cAIC < 2 as well as only for ADAS-cog, model (34) showed a Delta cAIC < 2 and thus substantial evidence compared to the best fitting model. All of these models included the interaction terms of the time variable with WMNE, as well as the second order interaction terms of the time variable with WMNE and with AV45 as well as TAU. Additionally, the best fitting model for CDRSOB also included the interaction term of WMNE with the time variable and WMHV. For an overview of the best fitting models, please refer to table 2.

**Table 2:** Comparison of best fitting models. Cog Out, cognitive outcome variable. No. Par, number of estimates parameters. Log(L), log-likelihood. cAIC, conditional Akaike information criterion. ΔcAIC, difference of the cAIC of the model considered and the model with lowest cAIC. ωcAIC, cAIC, difference of the cAIC of the model considered and the model with lowest cAIC. ωcAIC, cAIC, Akaike weights. R^2^m, marginal coefficient of determination. ADAS-cog, AD assessment scale. CDRSOB, clinical dementia rating sum of boxes. MMSE, minimental state examination.

| Cog Out | Model | No. Par | log(L) | cAIC | ΔcAIC | ωcAIC | R<sup>2</sup>m |
| --- | --- | --- | --- | --- | --- | --- | --- |
| ADAS-cog | 10 | 19 | -1248.1 | 2388.5 | 0.3266 | .2803 | .4045 |
| ADAS-cog | 31 | 18 | -1248.6 | 2388.2 | 0 | .3300 | .4120 |
| ADAS-cog | 34 | 20 | -1247.9 | 2389.6 | 1.4286 | .1615 | .4053 |
| CDRSOB | 34 | 20 | -580.1 | 1089.5 | 0 | .6528 | .4333 |
| MMSE | 10 | 19 | -791.3 | 1523.3 | 0.2682 | .2177 | .4092 |
| MMSE | 31 | 18 | -791.5 | 1523.1 | 0 | .2489 | .4155 |

Significance testing of these terms yielded significant positive interactions of WMNE with the time variable for CDRSOB and MMSE (p=0.03848 and p=0.01138, respectively), significant positive interactions of WMNE with the time variable and AV45 for all cognitive outcome measures (ADAS-cog: p=0.00463, CDRSOB: p<0.00001, MMSE: p=0.03549), significant negative interactions of WMNE with the time variable and TAU for all cognitive outcome variables (ADAS-cog: p=0.00446, CDRSOB: p<0.01535, MMSE: p=0.00164) as well as a significant negative interaction of WMNE with the time variable and WMHV for CDRSOB (p=0.01433). Positive interactions were such that at higher WMNE, time was less negatively associated with resilience for the interaction of WMNE with the time variable, while for the interaction of WMNE with the time variable and AV45 they were such that at higher amounts of AV45, the positive interaction of WMNE with the time variable was further increased. In contrast, negative interactions were sucht that at higher amounts of TAU or WMHV, the interaction of WMNE with the time variable was decreased. Robust reestimation of models yielded coefficients of the same directionality and comparable magnitude. FDR correction at 5% did not lead to the rejection of any of the results. P-values reestimated using parametric bootstrapping were similar to those calculated by LRT. Variance inflation factors were below 2 for all predictors in the best fitting models. For an overview of estimated coefficients and respective statistics please refer to table 3.

**Table 3:** Estimates of model terms of interest. Cog Out, Cognitive outcome. Std Beta, standardized regression coefficient. Rob Std Beta, standardized regression coefficient of robust mixed effects regression. FDR, false discovery rate. ADAS-cog, AD assessment scale. CDRSOB, clinical dementia rating sum of boxes. MMSE, minimental state examination.WMNE, WMNE.

| Cog Out | Term of Interest | Std<br>Beta | Rob Std<br>Beta | Chi<sup>2</sup> | p-value | FDR<br>corrected<br>p-value | p-value<br>param.<br>Boot-<br>strapping |
| --- | --- | --- | --- | --- | --- | --- | --- |
| ADAS-cog | T * WMNE | -0.004 | -0.009 | 0.044 | 0.83338 | .83338 | .83500 |
| ADAS-cog | T * WMNE * AV45 | 0.088 | 0.066 | 8.020 | 0.00463* | .01157* | .00516* |
| ADAS-cog | T * WMNE * TAU | -0.074 | -0.057 | 8.088 | 0.00446* | .01157* | .00501* |
| CDRSOB | T * WMNE | 0.098 | 0.090 | 4.284 | 0.03848* | .04275* | .04042* |
| CDRSOB | T * WMNE * AV45 | 0.276 | 0.216 | 56.736 | <0.00001* | <.00001* | <.00001* |
| CDRSOB | T * WMNE * TAU | -0.069 | -0.073 | 5.876 | 0.01535* | .02193* | .01667* |
| CDRSOB | T * WMNE * WMHV | -0.058 | -0.046 | 5.997 | 0.01433* | .02193* | .01574* |
| MMSE | T * WMNE | 0.074 | 0.085 | 6.405 | 0.01138* | .02193* | .01229* |
| MMSE | T * WMNE * AV45 | 0.024 | 0.026 | 4.422 | 0.03549* | .04275* | .03794* |
| MMSE | T * WMNE * TAU | -0.030 | -0.026 | 9.914 | 0.00164* | .00820* | .00182* |

Evaluating the results of model estimation and significance testing in conjunction with the resilience framework described in the previous section leads to the following four results: First, as all the best models for each cognitive outcome measure contained significant interaction terms of WMNE with the time variable and pathology measures, general resilience to cognitive decline could not be demonstrated. Second, dynamic resilience to cognitive decline was demonstrated for AV45 in all cognitive outcome measures considered due to the significant respective interaction terms. Third, limited resilience to cognitive decline was demonstrated for TAU in all cognitive outcome measures considered as well as for WMHV in CDRSOB due to the significant respective interaction terms. Fourth, the significant positive interaction terms of WMNE with the time variable for CDRSOB and MMSE demonstrate that on average of the modulating (demeaned) pathology measures in the respective models (AV45, TAU and WMHV for CDRSOB; AV45 and TAU for MMSE) WMNE is associated with greater resilience to cognitive decline for these cognitive outcome measures.

For a visualization of the estimated regression planes as well as scatter plots of the data points please refer to figures 1 and 2 as well as figures 3 and 4, respectively.

**Figure 1:**
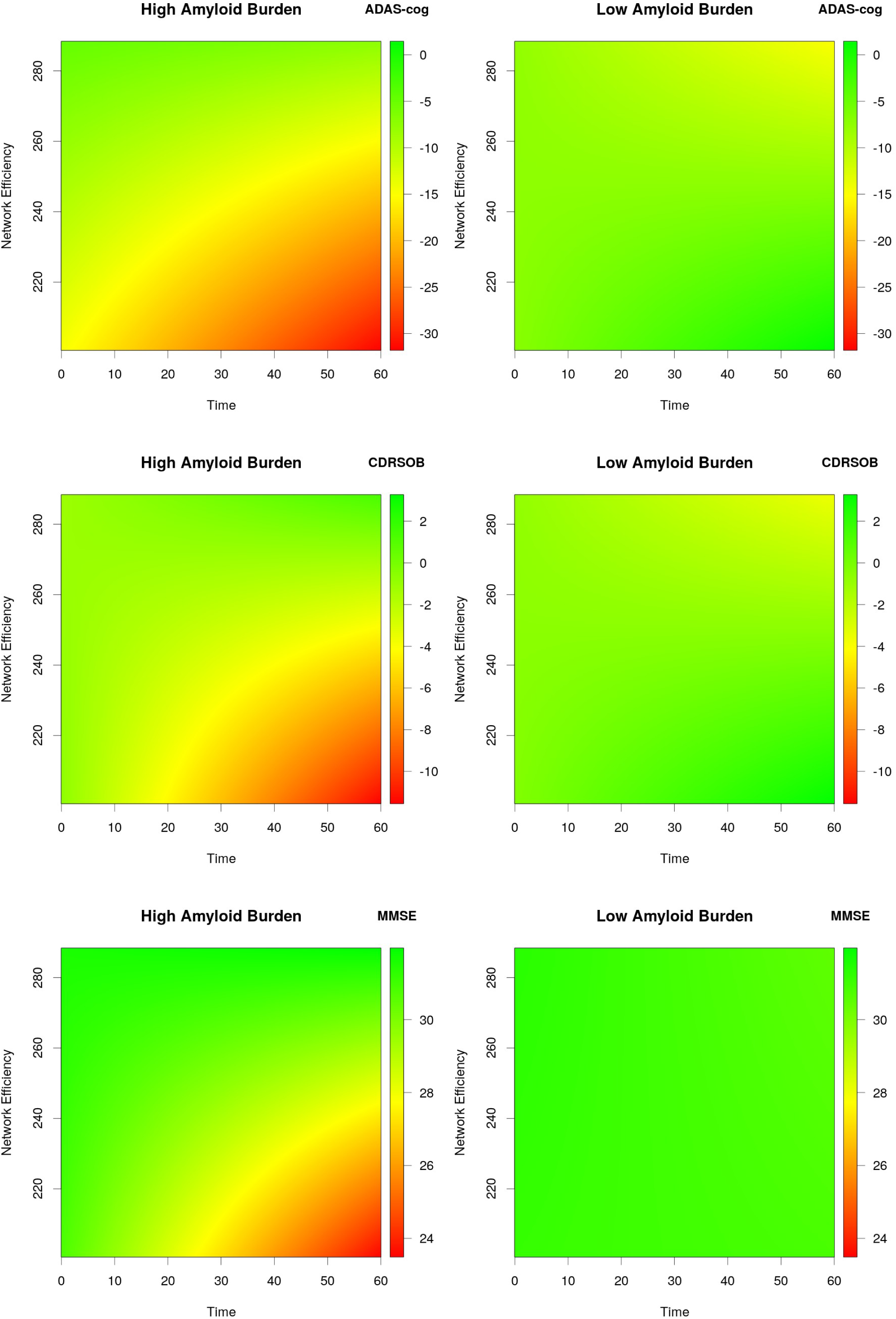
Dynamic resilience to cognitive decline at high amyloid burden – model estimates. The plots show cognitive outcome predicted by time (in months after baseline), white matter network efficiency and amyloid burden as well as their first and the second order interaction terms according to the estimated models, independent of other baseline covariates and pathology measures at mean +1 * standard deviation (high) and mean −1 * standard deviation (low) of amyloid burden. ADAS-cog, Alzheimer’s disease assessment scale. CDRSOB, clinical dementia rating sum of boxes. MMSE, minimental state examination.

**Figure 2:**
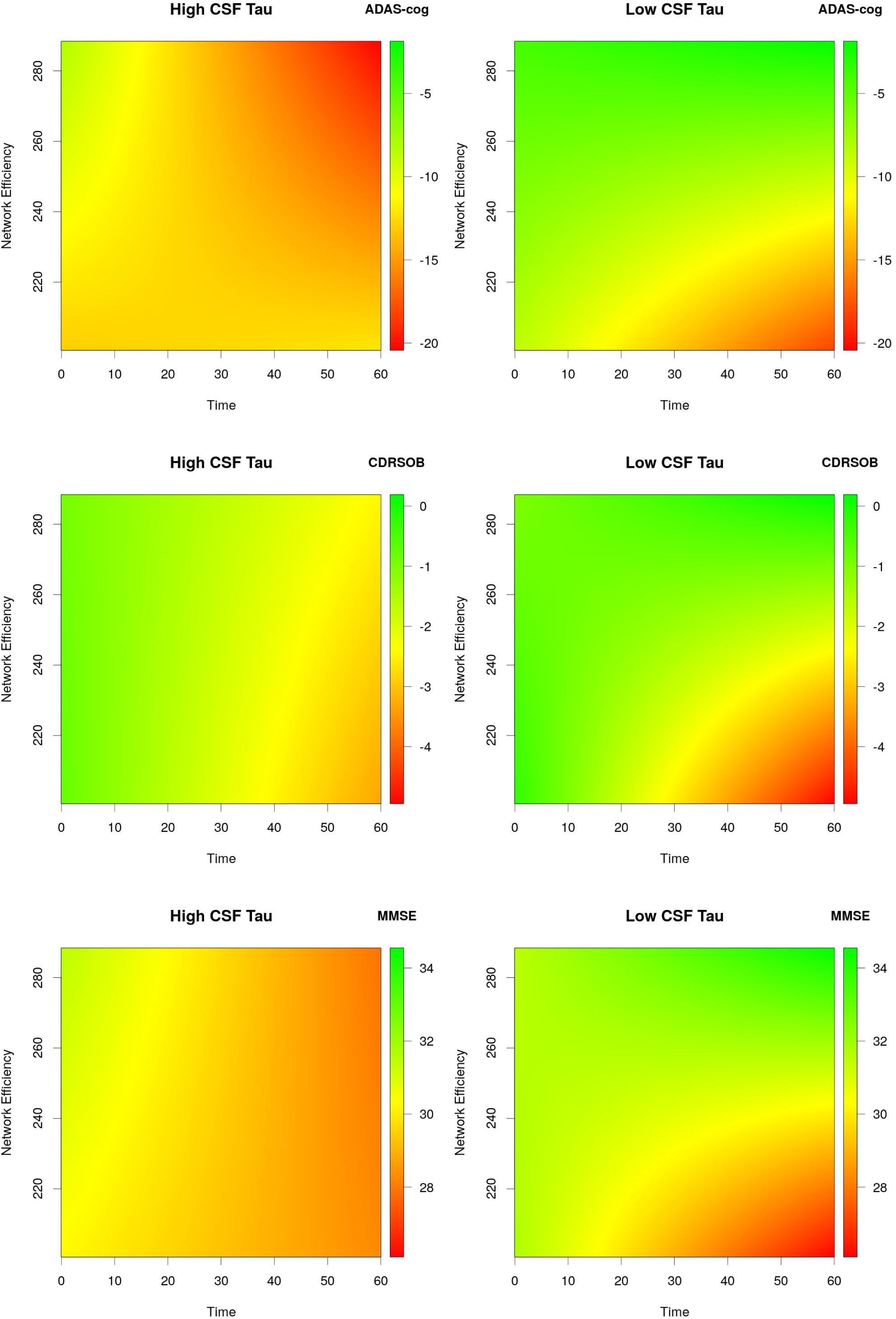
Limited general resilience to cognitive decline at low CSF tau – model estimates. The plots show cognitive outcome predicted by time (in months after baseline), white matter network efficiency and CSF tau as well as their first and the second order interaction terms according to the estimated models, independent of other baseline covariates and pathology measures at mean +1 * standard deviation (high) and mean −1 * standard deviation (low) of CSF tau. ADAS-cog, Alzheimer’s disease assessment scale. CDRSOB, clinical dementia rating sum of boxes. MMSE, minimental state examination.

**Figure 3:**
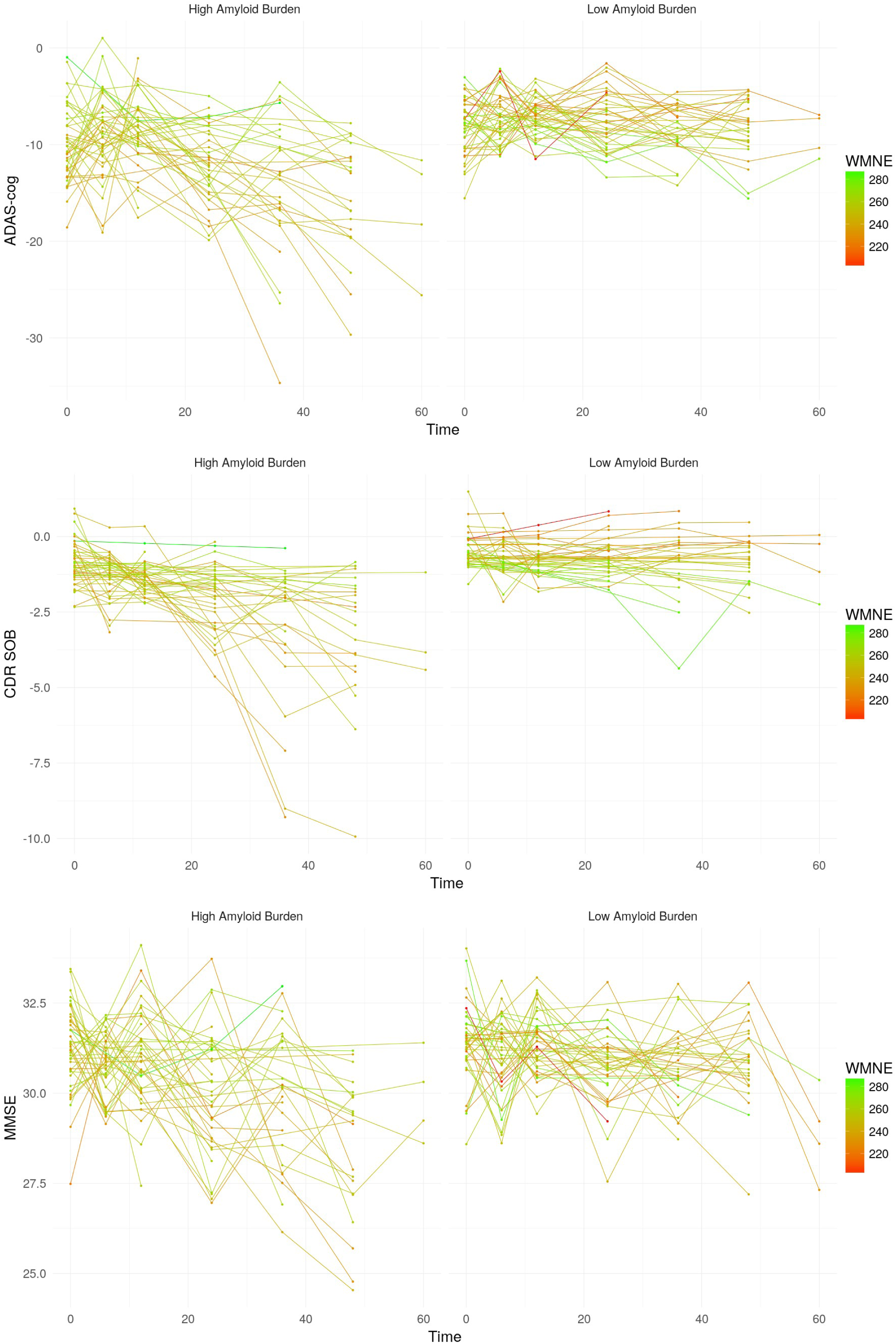
Dynamic resilience to cognitive decline at high amyloid burden – scatter plots. High amyloid burden vs. low amyloid burden, median split. WMNE, white matter network efficiency. Time, time in months after baseline. ADAS-cog, Alzheimer’s disease assessment scale. CDRSOB, clinical dementia rating sum of boxes. MMSE, minimental state examination.

**Figure 4:**
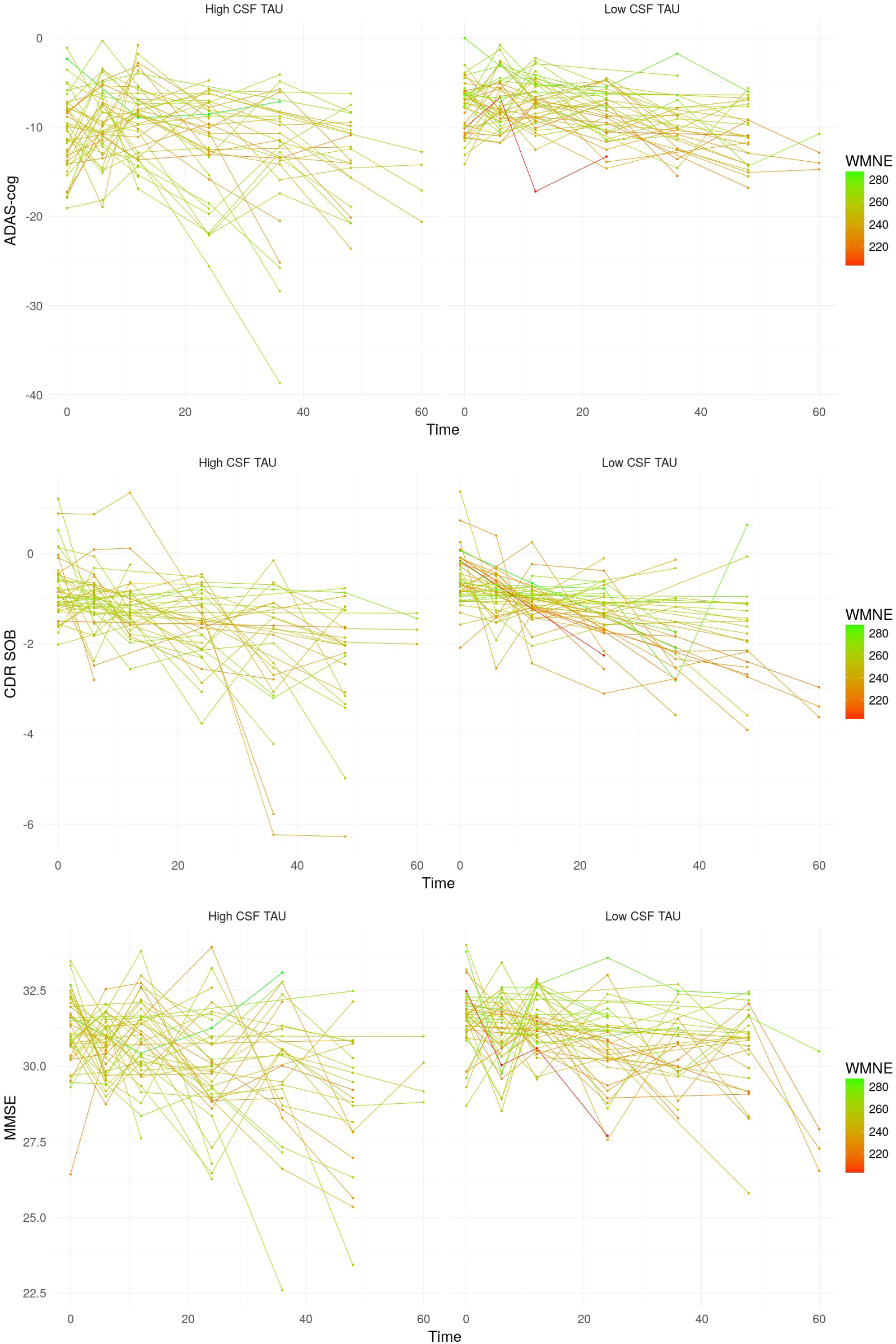
Limited general resilience to cognitive decline at low CSF tau – scatter plots. High CSF tau vs. low CSF tau, median split. WMNE, white matter network efficiency. Time, time in months after baseline. ADAS-cog, Alzheimer’s disease assessment scale. CDRSOB, clinical dementia rating sum of boxes. MMSE, minimental state examination.

## Discussion

The main results of this study indicate that in a population of non-demented elderly with varying amounts of age- and AD-related cerebral pathology, higher efficiency of the cerebreal network may be associated with more resilience to cognitive decline. This association was increased at higher amounts of cerebral amyloid burden and decreased at higher levels of CSF tau. WMNE may therefore be a factor of dynamic resilience to cognitive decline with respect to amyloid load whilst being a limited resilience factor with respect to tau burden.

To our best knownledge, this is the first study investigating the association of WM network properties and resilience to cognitive decline, quantified as lower cognitive decline unexplained by baseline cerebral pathology. However, other studies have demonstrated associations between WM network properties and other non-physiological resilience factors. Specifically, three studies demonstrated an association between intelligence and WMNE (Bathelt et al., 2018; Fischer et al., 2014; Li et al., 2009). A fourth study demonstrated an association between education and network flow, a measure quantifying hypothetical rerouting capabilities of the network(Wook Yoo et al., 2015). Although these studies are of limited comparability due to differences in methodology and sample composition, one may be tempted to speculate that WM network properties and WMNE specifically may form part of the physiological basis of intelligence and education as resilience factors. However, as education was controlled for in all analyses of the present study, the association of WMNE with resilience to cognitive decline seems to go beyond the effects of education.

A hypothetical mechanism by which WMNE may be associated with resilience to cognitive decline can be derived from its association with intelligence. Intelligence ratings consist of a number of cognitively demanding tests (Deary, Penke, & Johnson, 2010). The synchronized processing of these tasks by functional networks of distributed regions requires efficient and effective information transfer between them, which in turn depends on the structural connectivity and integrity of WM tracts (Penke et al., 2012). Additionally, there is evidence demonstrating that maintained functional connectivity is more associated with cognitive performance at higher age (Tsvetanov et al., 2016, 2018) and that the brain engages in compensatory activity in aging and the presence of cerebral neuropathology by recruiting additional neural resources across the brain (Reuter-Lorenz & Park, 2014; Stargardt et al., 2015), which probably likewise depends on structural integrity and efficient organization as a modulator of functional reorganization. Assuming that the brains compensatory reaction to age-associated pathological changes is alike to the compensatory reaction to aging, the findings of the present study could be explained as follows: as the aging brain accumulates deteriorative changes such as loss of grey matter volume and in some cases AD-related pathology such as amyloid accumulation, it engages compensatory processes aimed at recruiting more neural resources, whose recruitment for task processing depends upon information transfer efficiency, a surrogate measure of which may be WMNE. It follows that as the brain accumulates age- and, potentially, AD-related deteriorative changes with the passage of time relative to baseline, WMNE will become more important for sustained cognition and thus more associated with resilience. This is supported by the finding of significant positive interaction terms of the time variable with baseline WMNE estimated for two out of three cognitive outcome measures considered.

Interestingly, the association of WMNE with resilience to cognitive decline described above was increased in subjects with higher baseline cerebral amyloid load, thus indicating resilience to cognitive decline that is dynamic with respect to amyloid burden. This result was consistent across all cognitive outcome measures investigated and is also consistent with the resilience mechanism proposed above: elevated cerebral amyloid load has been demonstrated to be associated with a higher rate of cognitive decline (Donohue et al., 2017). However, it is not locally associated with GM atrophy at the earliest preclinical stages of the AD trajectory (Karran, Mercken, & Strooper, 2011; Kljajevic, Grothe, Ewers, & Teipel, 2014). In this scenario, compensation by recruitment of additional neural resources via the WM network would be both necessary as well as effective, as the to be recruited neural resources in the form of grey matter regions still remain mostly intact. However, the opposite may be true for increased levels of CSF tau at baseline, where the decrease of cognitive decline due to higher baseline WMNE was consistently lower for all three measures of cognitive outcome considered, thus indicating resilience to cognitive decline that is limited with respect to tau. Increased tau is usually accompanied by more clinically relevant cognitive impairment and neurodegeneration (Amlien et al., 2013; Solé-Padullés et al., 2011), especially at later stages of the AD-typical trajectory commonly referred to as the amyloid cascade (Jack et al., 2013). In this scenario, the to be recruited additional neural resources may already be impaired, which would probably render the compensatory functional reorganization less effective.

An alternative or perhaps complimentary explanation of the limited resilience to cognitive decline at increased levels of baseline CSF-tau is provided by the finding of WM microstructural integrity deterioration at increased levels of CSF-tau (Amlien et al., 2013). These may not be reflected in the WMNE measure considered in this study, but impair network function such that compensation via recruitment of additional neural resources is rendered less effective. The same explanation could be applicable to the finding of limited resilience to cognitive decline at higher baseline volume of WMHV, which are associated with deterioration in local WM microstructural integrity and cognition (Vernooij et al., 2009).

When considering the points discussed, one ought to bear in mind that any variance of WMNE associated with AV45, TAU or WMHV was partialled out for the estimation of all other model coefficients. This may seem contradictory to the points made in the previous paragraph, where it was argued that TAU and WMHV may deterioratively affect the network and thus impair its provision of compensatory capabilities, which seems likely at least for elevated WMHV. This argument implies that the putative effects of tau and WMHV on the network are reflected in the WMNE measure. If this had been the case, however, neither tau nor WMHV would have modulated the association of WMNE with resilience to cognitive decline. In order to improve WMNE as a surrogate measure of the network’s potential to support resilience, future studies might investigate modifications of WMNE such that likely changes due to tau and WMHV as demonstrated previously (Amlien et al., 2013; Vernooij et al., 2009) will be reflected within a modified WMNE measure.

Finally, apart from the mechanistic considerations discussed above, we believe that the results of this study may hold potential clinical relevance. The inclusion of WMNE in the best models for all cognitive outcome measures increases the models’ conditional R^2^ by a considerable 12.5% for ADAS-cog, 15.7% for CDRSOB and 9.3% for MMSE, as can be demonstrated by comparing the best models to the respective reduced models with all terms containing WMNE removed (data not shown). This demonstrates that WM structural connectivity in general and WMNE in particular may increase the accuracy of predictions of cognitive decline in elderly persons at risk of cognitive decline due to increased amounts of biomarkers of AD-typical pathology. Furthermore, future studies might investigate the factors that determine individual WMNE independent of cerebral pathology with the goal to develop intervention strategies aimed at improving resilience in individuals.

This study has several limitations. First, WMNE and pathology measures were measured at baseline. As such, further longitudinal associations or interactions between them could not be taken into account statistically. This means that the modulation of the association of WMNE with resilience to cognitive decline by pathology measures might have another alternative explanation: a lower association of resilience to cognitive decline with WMNE at higher baseline pathology could also be explained by a time lagged deteriorative association between the pathology measure and WMNE, whereas in this view a higher association between resilience to cognitive decline with WMNE at higher baseline pathology could be explained by time lagged (compensatory) neuroplasticity of WMNE as a response to increase baseline pathology. Second, the sample considered potentially includes subjects with future sporadic or familial AD, and the MCI cohort has additionally been enriched for memory impairment. As such, the variables considered probably do not follow a distribution representative for the general population, which may limit the generalizability of results. However, cognitive status (CN and MCI) was controlled for as a random variable in statistical modelling. Third, global pathology measures were used. Fourth, data were acquired at different sites and scanners. However, including center as an additional random effect did not alter results (data not shown).

## Conclusion

Higher WMNE may be associated with lower cognitive decline in cognitively healthy elderly and patients of MCI, especially so at the earlier stages of the AD biomarker cascade characterized by increased amounts of baseline cerebral amyloid load and low amounts of baseline CSF tau. WMNE or may thus be surrogate markers for the physiological basis of resilience to cognitive decline and provide a direction for further research aimed at improving the prediction of cognitive decline of persons at risk of dementia as well as research investigating factors leading to higher individual WMNE with the goal of designing intervention strategies aiming to improve individual resilience to cognitive decline.

## Informed consent

All procedures followed were in accordance with the ethical standards of the responsible committee on human experimentation (institutional and national) and with the Helsinki Declaration of 1975, and the applicable revisions at the time of the investigation. Informed consent was obtained from all patients for being included in the study.

## Non-Animal Research

No animal studies were carried out by the authors for this article.

## Data availability

The data that support the findings of this study are available from the Alzheimer’s Disease Neuroimaging Initiative (ADNI). Restrictions apply to the availability of these data due to privacy or ethical restrictions. Data are available at http://adni.loni.usc.edu/ with the permission of the ADNI.

## Acknowledgements

Data collection and sharing for this project was funded by the Alzheimer’s Disease Neuroimaging Initiative (ADNI) (National Institutes of Health Grant U01 AG024904) and DOD ADNI (Department of Defense award number W81XWH-12-2-0012). ADNI is funded by the National Institute on Aging, the National Institute of Biomedical Imaging and Bioengineering, and through generous contributions from the following: AbbVie, Alzheimer’s Association; Alzheimer’s Drug Discovery Foundation; Araclon Biotech; BioClinica, Inc.; Biogen; Bristol-Myers Squibb Company; CereSpir, Inc.; Cogstate; Eisai Inc.; Elan Pharmaceuticals, Inc.; Eli Lilly and Company; EuroImmun; F. Hoffmann-La Roche Ltd and its affiliated company Genentech, Inc.; Fujirebio; GE Healthcare; IXICO Ltd.; Janssen Alzheimer Immunotherapy Research & Development, LLC.; Johnson & Johnson Pharmaceutical Research & Development LLC.; Lumosity; Lundbeck; Merck & Co., Inc.; Meso Scale Diagnostics, LLC.; NeuroRx Research; Neurotrack Technologies; Novartis Pharmaceuticals Corporation; Pfizer Inc.; Piramal Imaging; Servier; Takeda Pharmaceutical Company; and Transition Therapeutics. The Canadian Institutes of Health Research is providing funds to support ADNI clinical sites in Canada. Private sector contributions are facilitated by the Foundation for the National Institutes of Health (www.fnih.org). The grantee organization is the Northern California Institute for Research and Education, and the study is coordinated by the Alzheimer’s Therapeutic Research Institute at the University of Southern California. ADNI data are disseminated by the Laboratory for Neuro Imaging at the University of Southern California.

