## Supplement for "Network efficiency predicts resilience to cognitive decline in elderly at risk for Alzheimer’s"

**Appendix**

**Description of the statistical approach to investigate resilience**

Any analysis of resilience requires at least three types of measures: a possible resilience factor, a measure of pathology and a measure of cognitive outcome (Stern, 2012). However, analyses that include these three measures in regression analyses traditionally only consider a positive interaction between pathology and the possible resilience factor as evidence of resilience (Craik, Salthouse, & Salthouse, 2011). Recently, we published a reconceptualization of this approach that aims to operationalize a differentiation of resilience by its relation to pathology (Wolf et al., 2018). The investigation of resilience in the present paper is based on an extension of this approach.

Specifically, we model resilience as cognitive outcome not explained by pathology measures (see table 2 for an overview of pathology measures). A candidate resilience factor could then be associated with resilience in three ways. First, the association between resilience factor and resilience can be unmodulated by pathology measures, which requires that the mechanism behind the association of the resilience factor and resilience remains intact even in the presence of increased pathology. This is referred to as *general resilience*. Second, the association between resilience factor and resilience could be positively modulated by one or more pathology measures, indicating that the importance of the resilience factor for better cognition than predicted by pathology increases with increasing amounts of the respective pathology. This is termed *dynamical resilience*. Third, a resilience factor for which general or dynamic resilience has been established may additionally show an association with resilience that is negatively modulated by a pathology measure. This indicates that while the resilience factor is associated with resilience in certain scenarios – either on average with respect to the considered pathology measure or dynamically with respect to another pathology measure – the association with resilience diminishes with increasing amounts of the respective pathology measure. This is termed *limited resilience*.

These concepts can be statistically implemented using the following model equation:

COG ~ RESF + PATH_1_ + PATH_2_ + … + PATH_n_ + RES * (PATH_1_ + PATH_2_ + … +

PATH_n_)

where COG represents the cognitive outcome measure, PATH represents a measure of pathology, n is the number of pathology measures considered and RESF represents a possible resilience factor. Note that as PATH is partialled out of all other variables by the regression estimation, COG as well as RESF represent the variance of these measures unexplained by PATH. For this reason COG is referred to as resilience (Wolf et al., 2018).

This model supports the three types of resilience in the following cases:

**General resilience** – the term RESF is positively associated with COG, while none of the RESF * PATH terms are.

**Dynamic resilience** – one or more of the terms RESF * PATH are positively associated with COG. RESF can be positively associated with COG as well, in this case one could add that *on average* of PATH, RESF is positively associated with COG.

**Limited resilence** – either RESF and/or one or more of the terms RESF * PATH are positively associated with COG (i.e. general or dynamic resilience). In addition one or more of the remaining (i.e. different) terms RESF * PATH are negatively associated with COG.

Finally, the concepts of general and dynamic resilience overlap but are not equivalent to the concept of brain reserve and cognitive reserve put forward by(Barulli & Stern, 2013): whereas the definitions of brain reserve and cognitive reserve refer to the quality of the underlying mechanism of resilience, the concept of general and dynamic resilience is based entirely on the empirical manifestation of resilience as associations with the resilience factor(Wolf et al., 2018). Brain reserve could thus manifest as general resilience or dynamic resilience (e.g. dynamic in the case of a threshold based association), as might cognitive reserve (which may have a positive effect on cognition even in the absence of pathology).

**Model adaption to longitudinal data**

As the aim of the present study is to investigate resilience to cognitive decline, i.e. lower cognitive decline than expected based on baseline pathology measures, the base resilience model described in the previous section needs to be extended. The original model (with all pathology measures collapsed into PATH for readability) is given as

  	COG ~ RESF * PATH + RESF + PATH.

To accommodate longitudinal data and model the resilience to cognitive decline, this model is extended by the addition of a time variable T as well as its interaction terms with the other independent variables:

COG ~ (RESF * PATH + RESF + PATH) * T + T

Note that the effect of PATH as well as the modulation of the association of the T with COG by PATH, i.e. cognitive decline due to baseline pathology, are partialled out of COG. In analogy to the original model, this extended model supports resilience to cognitive decline in the following scenarios:

**General resilience to cognitive decline** – the term RESF * T is positively associated with COG, while none of the RESF * T * PATH terms are.

**Dynamic resilience to cognitive decline** – one or more of the terms RESF * T * PATH are positively associated with COG. RESF * T can be positively associated with COG as well, in this case one could add that *on average* of PATH, RESF * T is associated with COG.

**Limited resilence to cognitive decline** – either RESF * T and/or one or more of the terms RESF * T * PATH are positively associated with COG (i.e. general or dynamic resilience to cognitive decline). In addition one or more of the remaining (i.e. different) terms RESF * T * PATH are negatively associated with COG.

**List of the family of models considered and evaluated**

The following list generally only contains the present highest order interaction terms for brevity, their constituent terms were always included; WMNE: white matter network efficiency, WMHV: white matter hyperintensity volume.

```
 [1] "WMNE + T + AV45 + TAU + WMHV + Age + Gender + APOE4 + Education + WMNE*T + WMNE*AV45 + T*AV45 + WMNE*TAU + T*TAU + T*WMHV + WMNE*T*AV45  + (1|Subject) + (1|Diagnosis)"
 [2] "WMNE + T + AV45 + WMHV + TAU + Age + Gender + APOE4 + Education + WMNE*T + WMNE*AV45 + T*AV45 + WMNE*WMHV + T*TAU + T*WMHV + WMNE*T*AV45  + (1|Subject) + (1|Diagnosis)"
 [3] "WMNE + T + AV45 + TAU + WMHV + Age + Gender + APOE4 + Education + WMNE*T + WMNE*AV45 + T*AV45 + WMNE*TAU + WMNE*WMHV + T*TAU + T*WMHV + WMNE*T*AV45  + (1|Subject) + (1|Diagnosis)"
 [4] "WMNE + T + TAU + AV45 + WMHV + Age + Gender + APOE4 + Education + WMNE*T + WMNE*TAU + T*TAU + WMNE*AV45 + T*AV45 + T*WMHV + WMNE*T*TAU  + (1|Subject) + (1|Diagnosis)"
 [5] "WMNE + T + TAU + WMHV + AV45 + Age + Gender + APOE4 + Education + WMNE*T + WMNE*TAU + T*TAU + WMNE*WMHV + T*AV45 + T*WMHV + WMNE*T*TAU  + (1|Subject) + (1|Diagnosis)"
 [6] "WMNE + T + TAU + AV45 + WMHV + Age + Gender + APOE4 + Education + WMNE*T + WMNE*TAU + T*TAU + WMNE*AV45 + WMNE*WMHV + T*AV45 + T*WMHV + WMNE*T*TAU  + (1|Subject) + (1|Diagnosis)"
 [7] "WMNE + T + WMHV + AV45 + TAU + Age + Gender + APOE4 + Education + WMNE*T + WMNE*WMHV + T*WMHV + WMNE*AV45 + T*AV45 + T*TAU + WMNE*T*WMHV  + (1|Subject) + (1|Diagnosis)"
 [8] "WMNE + T + WMHV + TAU + AV45 + Age + Gender + APOE4 + Education + WMNE*T + WMNE*WMHV + T*WMHV + WMNE*TAU + T*AV45 + T*TAU + WMNE*T*WMHV  + (1|Subject) + (1|Diagnosis)"
 [9] "WMNE + T + WMHV + AV45 + TAU + Age + Gender + APOE4 + Education + WMNE*T + WMNE*WMHV + T*WMHV + WMNE*AV45 + WMNE*TAU + T*AV45 + T*TAU + WMNE*T*WMHV  + (1|Subject) + (1|Diagnosis)"
[10] "WMNE + T + AV45 + TAU + WMHV + Age + Gender + APOE4 + Education + WMNE*T + WMNE*AV45 + T*AV45 + WMNE*TAU + T*TAU + WMNE*WMHV + T*WMHV + WMNE*T*AV45 + WMNE*T*TAU  + (1|Subject) + (1|Diagnosis)"
[11] "WMNE + T + AV45 + WMHV + TAU + Age + Gender + APOE4 + Education + WMNE*T + WMNE*AV45 + T*AV45 + WMNE*WMHV + T*WMHV + WMNE*TAU + T*TAU + WMNE*T*AV45 + WMNE*T*WMHV  + (1|Subject) + (1|Diagnosis)"
[12] "WMNE + T + TAU + WMHV + AV45 + Age + Gender + APOE4 + Education + WMNE*T + WMNE*TAU + T*TAU + WMNE*WMHV + T*WMHV + WMNE*AV45 + T*AV45 + WMNE*T*TAU + WMNE*T*WMHV  + (1|Subject) + (1|Diagnosis)"
[13] "WMNE + AV45 + T + TAU + WMHV + Age + Gender + APOE4 + Education + WMNE*AV45 + AV45*T + T*TAU + T*WMHV  + (1|Subject) + (1|Diagnosis)"
[14] "WMNE + TAU + T + AV45 + WMHV + Age + Gender + APOE4 + Education + WMNE*TAU + T*AV45 + TAU*T + T*WMHV  + (1|Subject) + (1|Diagnosis)"
[15] "WMNE + T + AV45 + TAU + WMHV + Age + Gender + APOE4 + Education + WMNE*T + T*AV45 + T*TAU + T*WMHV  + (1|Subject) + (1|Diagnosis)"
[16] "WMNE + WMHV + T + AV45 + TAU + Age + Gender + APOE4 + Education + WMNE*WMHV + T*AV45 + T*TAU + WMHV*T  + (1|Subject) + (1|Diagnosis)"
[17] "WMNE + AV45 + TAU + T + WMHV + Age + Gender + APOE4 + Education + WMNE*AV45 + WMNE*TAU + AV45*T + TAU*T + T*WMHV  + (1|Subject) + (1|Diagnosis)"
[18] "WMNE + AV45 + T + TAU + WMHV + Age + Gender + APOE4 + Education + WMNE*AV45 + WMNE*T + AV45*T + T*TAU + T*WMHV  + (1|Subject) + (1|Diagnosis)"
[19] "WMNE + AV45 + WMHV + T + TAU + Age + Gender + APOE4 + Education + WMNE*AV45 + WMNE*WMHV + AV45*T + T*TAU + WMHV*T  + (1|Subject) + (1|Diagnosis)"
[20] "WMNE + TAU + T + AV45 + WMHV + Age + Gender + APOE4 + Education + WMNE*TAU + WMNE*T + T*AV45 + TAU*T + T*WMHV  + (1|Subject) + (1|Diagnosis)"
[21] "WMNE + TAU + WMHV + T + AV45 + Age + Gender + APOE4 + Education + WMNE*TAU + WMNE*WMHV + T*AV45 + TAU*T + WMHV*T  + (1|Subject) + (1|Diagnosis)"
[22] "WMNE + T + WMHV + AV45 + TAU + Age + Gender + APOE4 + Education + WMNE*T + WMNE*WMHV + T*AV45 + T*TAU + T*WMHV  + (1|Subject) + (1|Diagnosis)"
[23] "WMNE + AV45 + TAU + T + WMHV + Age + Gender + APOE4 + Education + WMNE*AV45 + WMNE*TAU + WMNE*T + AV45*T + TAU*T + T*WMHV  + (1|Subject) + (1|Diagnosis)"
[24] "WMNE + AV45 + TAU + WMHV + T + Age + Gender + APOE4 + Education + WMNE*AV45 + WMNE*TAU + WMNE*WMHV + AV45*T + TAU*T + WMHV*T  + (1|Subject) + (1|Diagnosis)"
[25] "WMNE + AV45 + T + WMHV + TAU + Age + Gender + APOE4 + Education + WMNE*AV45 + WMNE*T + WMNE*WMHV + AV45*T + T*TAU + T*WMHV  + (1|Subject) + (1|Diagnosis)"
[26] "WMNE + TAU + T + WMHV + AV45 + Age + Gender + APOE4 + Education + WMNE*TAU + WMNE*T + WMNE*WMHV + T*AV45 + TAU*T + T*WMHV  + (1|Subject) + (1|Diagnosis)"
[27] "WMNE + AV45 + TAU + T + WMHV + Age + Gender + APOE4 + Education + WMNE*AV45 + WMNE*TAU + WMNE*T + WMNE*WMHV + AV45*T + TAU*T + T*WMHV  + (1|Subject) + (1|Diagnosis)"
[28] "WMNE + T + AV45 + TAU + WMHV + Age + Gender + APOE4 + Education + WMNE*T + WMNE*AV45 + T*AV45 + T*TAU + T*WMHV + WMNE*T*AV45  + (1|Subject) + (1|Diagnosis)"
[29] "WMNE + T + TAU + AV45 + WMHV + Age + Gender + APOE4 + Education + WMNE*T + WMNE*TAU + T*TAU + T*AV45 + T*WMHV + WMNE*T*TAU  + (1|Subject) + (1|Diagnosis)"
[30] "WMNE + T + WMHV + AV45 + TAU + Age + Gender + APOE4 + Education + WMNE*T + WMNE*WMHV + T*WMHV + T*AV45 + T*TAU + WMNE*T*WMHV  + (1|Subject) + (1|Diagnosis)"
[31] "WMNE + T + AV45 + TAU + WMHV + Age + Gender + APOE4 + Education + WMNE*T + WMNE*AV45 + T*AV45 + WMNE*TAU + T*TAU + T*WMHV + WMNE*T*AV45 + WMNE*T*TAU  + (1|Subject) + (1|Diagnosis)"
[32] "WMNE + T + AV45 + WMHV + TAU + Age + Gender + APOE4 + Education + WMNE*T + WMNE*AV45 + T*AV45 + WMNE*WMHV + T*WMHV + T*TAU + WMNE*T*AV45 + WMNE*T*WMHV  + (1|Subject) + (1|Diagnosis)"
[33] "WMNE + T + TAU + WMHV + AV45 + Age + Gender + APOE4 + Education + WMNE*T + WMNE*TAU + T*TAU + WMNE*WMHV + T*WMHV + T*AV45 + WMNE*T*TAU + WMNE*T*WMHV  + (1|Subject) + (1|Diagnosis)"
[34] "WMNE + T + AV45 + TAU + WMHV + Age + Gender + APOE4 + Education + WMNE*T + WMNE*AV45 + T*AV45 + WMNE*TAU + T*TAU + WMNE*WMHV + T*WMHV + WMNE*T*AV45 + WMNE*T*TAU + WMNE*T*WMHV  + (1|Subject) + (1|Diagnosis)"
[35] "WMNE + T + AV45 + TAU + WMHV + Age + Gender + APOE4 + Education + T*AV45 + T*TAU + T*WMHV  + (1|Subject) + (1|Diagnosis)"
[36] "T + AV45 + TAU + WMHV + Age + Gender + APOE4 + Education + T*AV45 + T*TAU + T*WMHV  + (1|Subject) + (1|Diagnosis)"
```

Barulli, D., & Stern, Y. (2013). Efficiency, capacity, compensation, maintenance, plasticity: emerging concepts in cognitive reserve. *Trends in Cognitive Sciences*, *17*(10), 502–509.

Craik, F. I. M., Salthouse, T. A., & Salthouse, T. A. (2011, March 15). Intelligence, Education, and the Brain Reserve Hypothesis: Helen Christensen, Kaarin J. Anstey, Liana S. Leach, and Andrew J. Mackinnon. *The Handbook of Aging and Cognition*. Retrieved September 20, 2018, from https://www.taylorfrancis.com/

Stern, Y. (2012). Cognitive reserve in ageing and Alzheimer’s disease. *The Lancet Neurology*, *11*(11), 1006–1012.

Wolf, D., Fischer, F. U., & Fellgiebel, A. (2018). A methodological approach to studying resilience mechanisms: demonstration of utility in age and Alzheimer’s disease-related brain pathology. *Brain Imaging and Behavior*, 1–10.
